# Cooperative nest building in wild jackdaw pairs

**DOI:** 10.1101/2020.12.15.422933

**Authors:** Luca G. Hahn, Rebecca Hooper, Guillam E. McIvor, Alex Thornton

## Abstract

Animals create diverse structures, both individually and cooperatively, using materials from their environment. One striking example are the nests birds build for reproduction, which protect the offspring from external stressors such as predators and temperature, promoting reproductive success. To construct a nest successfully, birds need to make various decisions, for example regarding the nest material and their time budgets. To date, research has focused mainly on species where one sex is primarily responsible for building the nest. In contrast, the cooperative strategies of monogamous species in which both sexes contribute to nest building are poorly understood. Here we investigated the role of both sexes in nest building and fitness correlates of behaviour in wild, monogamous jackdaw pairs (*Corvus monedula*). We show that both partners contributed to nest building and behaved similarly, with females and males present in the nest box for a comparable duration and transporting material to the nest equally often. However, while females spent more time constructing the nest, males tended to invest more time in vigilance, potentially as a means of coping with competition for nest cavities. These findings suggest a moderate degree of division of labour, which may facilitate cooperation. Moreover, some aspects of behaviour were related to proxies of reproductive success (lay date and egg volume). Females that contributed relatively more to bringing material laid earlier clutches and pairs that spent less time together in the nest box had larger eggs. Thus, selection pressures may act on how nest building pairs spend their time and cooperatively divide the labour. We conclude that cooperative nest building in birds could be associated with monogamy and obligate biparental care, and provides a vital but relatively untapped context through which to study the evolution of cooperation.

**Highlights:**

- In wild monogamous jackdaws, mates behaved similarly and cooperated to build their nest.
- Females built more and called more frequently; males tended to be more vigilant.
- Females that contributed relatively more to transporting nest material laid earlier clutches.
- Pairs that spent more time together in the nest box had smaller eggs.
- Cooperation may be crucial in light of obligate biparental care and nest site competition.

## 1. Introduction

Across the animal kingdom, species build structures for various purposes relevant for fitness. Such animal architecture (Hansell, 2005, 2007) is used in diverse contexts such as creating a protective shelter (Rosell et al., 2005), reproduction and parental care (Deeming & Reynolds, 2015), capture of prey (Hunt, 1996), and communication and signalling (Borgia, 1995). A striking example are bird nests built for reproduction (Collias, 1964; Collias & Collias, 1984; Hansell, 2000; Healy, Walsh, & Hansell, 2008), which influence fitness by protecting the offspring, for example from predators through camouflage (Bailey et al., 2015) and from environmental stressors, such as temperature (Collias, 1964). Additionally, nests can function as an intraspecific signal of investment in reproduction (Massoni et al., 2012; Soler et al., 1998) and to attract mates (Metz et al., 2009). While nest building behaviour was traditionally assumed to be genetically predetermined (Nickell, 1958), recent evidence highlights an important role for learning (Bailey et al., 2014; Breen et al., 2016; Walsh et al., 2013). For example, male zebra finches (*Taeniopygia guttata*) adjust their preferred material based on their success in a past breeding attempt (Muth & Healy, 2011). However, research to date has focussed on species in which single individuals (often males) predominantly build the nest: in zebra finches, for instance, studies have focused on males, who are responsible for bringing the nest material (Zann, 1996). While both partners may then contribute to arranging the material in the nest, their cooperative interactions at this stage have not been investigated in detail. While there has been some work describing contributions to nest building in cooperative breeders like sociable weavers (*philetairus socius*) and white-browed sparrow-weavers (*Plocepasser mahali*) (Collias & Collias, 2008; Collias & Collias, 1978) cooperative nest building by monogamous mates remains largely unexplored. This is particularly surprising given that monogamy and biparental care are common in the majority of bird species (Black, 1996; Reichard & Boesch, 2013). There is therefore a need to investigate whether and how monogamous birds cooperate during nest building. This will allow us to comprehensively understand the costs and benefits of cooperation between partners during this key stage of the breeding cycle, and, more broadly, will allow a deeper insight into the cooperative behaviours underlying animal architecture.

Effective cooperation between mates can be vital for fitness, particularly in species with obligate biparental care (Griffith, 2019). However, the interests of both sexes do not align exactly, generating sexual conflict (Chapman et al., 2003; Harrison et al., 2009). Research has concentrated largely on how conflicts between mates are resolved when provisioning food to offspring (Hinde & Kilner, 2007; Iserbyt et al., 2015; Johnstone et al., 2014), making monogamous birds central study systems to understand the evolution of cooperative strategies. For instance, theoretical and empirical studies suggest that forms of conditional cooperation, such as turn-taking, may serve to reduce conflicts of interest and stabilise cooperation (Johnstone & Savage, 2019; Savage et al., 2017). Given that monogamous birds have long served as important model systems for understanding the evolution of cooperation, and that mates in some species are known to build the nest together (Birkhead, 2010; Massoni et al., 2012), it is striking that cooperative nest building strategies have rarely been examined explicitly. Establishing the role of the two sexes during cooperative nest building is crucial to our understanding of both cooperative strategies and animal architecture.

In birds, the degree of cooperation between the sexes during nest building could be linked to the mating system. For instance, in various polygynous weaver species (Ploceidae) males build nests alone to attract females (Bailey et al., 2016), whereas in monogamous weavers mated pairs build their nest cooperatively (Habig, 2020). Furthermore, two largely genetically monogamous species, Eurasian magpies (*Pica pica*) (Parrot, 1995) and rufous horneros (*Furnarius rufus*) (Diniz et al., 2019), also build their nest cooperatively (Birkhead, 2010; Massoni et al., 2012). However, fine-scale behaviours and time budgets have not previously been explored, so cooperative nest building and its fitness consequences remain poorly understood. The degree to which partners cooperate is likely to depend on how much their interests align. In species showing obligate biparental care, mates should invest (relatively equally) in the offspring, because a lack of investment by either parent is likely to lead to failure of the reproductive attempt (Clutton-Brock, 2019). Moreover, one could expect greater degrees of cooperation in species with low rates of extra-pair fertilisation (Lv et al., 2019), and high paternity certainty (Disciullo et al., 2019) as these conditions create highly interdependent fitness outcomes. The success of a clutch could be impacted by how bird pairs cooperate during nest building because cooperation may influence nest quality and because this process is energetically and temporally costly and (Collias, 1964; Mainwaring & Hartley, 2013). The energetic costs of nest building could vary between sexes due to differences in morphology, physiology, energetic demands, and available information. Consequently, while both mates may behave similarly, sexbased differences in the costs associated with certain activities could promote task specialisation, as shown by evolutionary individual-based simulations of individuals providing two types of parental care associated with a sex-based asymmetry regarding the costs (Barta et al., 2014). This could be important in the context of nest building as well; for example, male magpies and female rufous horneros bring relatively more material to the nest than the opposite sex. Investigating the roles of sexes, the level of cooperation, whether cooperation is repeatable within pairs, and the fitness consequences during nest building is also vital to further understand how individuals cope with the informational demands of decision-making processes whilst tracking another individual’s behaviour (Emery et al., 2007). Tracking each other’s behaviour could favour greater levels of behavioural synchrony, which could also be related to behavioural compatibility between partners, potentially resulting in more effective cooperation and greater reproductive success (Spoon et al., 2006).

Jackdaws (*Corvus monedula*) provide a particularly suitable study system to investigate cooperation during nest building. They are a highly social, colony-breeding corvid that forms long-term pair bonds (Lorenz, 1931; Wechsler, 1989). Pairs produce one clutch per year, with both sexes providing care to altricial chicks (Henderson & Hart, 1993). Moreover, unlike most socially monogamous bird species, jackdaws are genetically monogamous, so the reproductive success of partners is more interdependent than in species where extra-pair offspring occur (Gill et al., 2020). For jackdaws to reproduce successfully, they must build a nest before egg laying commences. Jackdaws build nests within cavities, and these consist of a platform (made of sticks) and a cup with soft material. Tightly linked fitness outcomes may generate selection pressure for cooperation between partners throughout the breeding season, including during the nest building stage.

This study had two main objectives: **(i)** To quantify behaviours and time budgets of pairs. We hypothesised that birds should cooperate and share the workload to build the nest, because both individuals should benefit from dividing costs and from a high-quality nest. Firstly, we predicted females and males should behave similarly by investing in the nest directly (e.g. by bringing nest material) and indirectly (e.g. through vigilance) (prediction 1). Secondly, however, we predicted that the time invested in these behaviours may not be symmetrical between the sexes given morphological, physiological, and informational differences (prediction 2). **(ii)** To examine the ultimate function of behaviours during nest building by investigating three different fitness proxies: relative lay date, clutch size, and egg volume. Laying earlier clutches can be advantageous and is often linked to reproductive success in birds (Perrins, 1965, 1970; Goumas et al., in prep.), for example because earlier layers face less competition in finding food for their young. Larger eggs could potentially provide the embryo with more resources, aiding its development and increasing the probability to survive (Krist, 2011). We hypothesised that how much birds invest in their nest and how they share the workload could be associated with reproductive success, with pairs that invest more overall and divide the labour (so that males contribute at least equally) being favoured. While females should invest substantially in the nest because they may be better informed about their requirements for incubation, males should contribute equally because this may allow females to invest more resources in the clutch, potentially maximising reproductive success. Furthermore, investment may determine the time to build the nest as well as its quality, which in turn could enhance embryo development and survival. Firstly, we predicted that how pairs allocate their time between different activities could impact fitness. More specifically, pairs investing more in the nest relative to other activities, such as vigilance, should lay earlier, and have larger clutches and eggs (prediction 3). Moreover, pairs that show greater total investment in the nest should lay earlier clutches (prediction 4). Pairs in which males invest at least equally in the nest as well as in nest site defence should lay earlier and produce larger clutches and eggs (prediction 5). If the optimal solution was for both individuals to invest equally, one might expect a quadratic relationship between relative contributions of females compared to the overall investment and fitness proxies. Finally, we predicted that jackdaws behaving more synchronously by spending more time together in the nest box should lay earlier and have larger clutches and eggs (prediction 6). As selection on nest building behaviours may depend on the degree to which they constitute repeatable traits, we also investigated the repeatability of behaviour over time.

## 2. Methods

### Ethics Statement

This study was conducted with approval from the University of Exeter Biosciences Research Ethics Committee (eCORN002970), following the ASAB Guidelines for the Treatment of Animals in Behavioural Research (ASAB, 2012). Jackdaws had previously been colour ringed for individual identification by qualified ringers licenced by the British Trust for Ornithology. The sex of each individual was confirmed through molecular sexing of blood samples (Griffiths, Double, Orr, & Dawson, 1998) under a UK Home Office licence (project licence 30/3261). Morphometrics of individuals, such as wing length, tarsus length, and body weight, were measured when temporarily capturing birds for ringing (see Greggor et al., 2017 for details). We used the exact age if birds were ringed as nestlings or juveniles, and for adults that were already present when field sites were established in 2012, we assigned the maximum age of eight years.

### Data collection

As cavity nesters, jackdaws accept nest boxes and typically return to the same nest site across years, allowing researchers to monitor behaviour by fitting boxes with video cameras. We recorded nest building behaviour of free-living jackdaws using CCTV cameras with integrated microphones hidden inside nest boxes during the breeding seasons of 2018 and 2019 at three breeding colonies in Cornwall, UK: X (50°10’23”N; 5°7’12”W), Y (50°11’26”N, 5°10’51”W), and Z (50°11’56”N, 5°10’9”W). Nest boxes were filmed slightly but significantly closer to the lay date in 2019 (mean ± SD = 7.69 ± 5.36 days) than in 2018 (10.83 ± 6.41 days) (LM, β ± SE = – 3.136 ± 1.551, t_59_ = – 2.022, *P* = 0.048). In total, we recorded 183.04 hours of video data (N = 62 videos; one video = one observation; mean video length ± SD 2.95 ± 1.07, range = 1 – 5 hours) from 40 distinct, breeding jackdaw pairs across 40 different nest boxes (N = 5 videos in 5 boxes at colony X; 27 videos in 15 boxes at Y; and 30 videos in 20 boxes at Z). All observations were conducted in the morning (start time 0630 – 0930 hours) to minimise the confounding effect of changing behavioural patterns throughout the day. We recorded a minimum of one video at each nest during the middle of the nest building phase in April. In seven cases, we recorded more than one video per pair per breeding season, and for 14 pairs, we recorded at least one video in each of 2018 and 2019. Boxes were monitored regularly throughout the breeding season to document lay date, clutch size, and egg volume. Measures of egg length and width served to calculate egg volume (Troscianko, 2014). In all observations, jackdaws built a nest, and all but one pair (box Z28, 2018), which was displaced by another pair, laid eggs.

### Video analysis

We analysed videos in a randomised order with regards to ‘year’ and ‘study site’, using the software *BORIS* version 7.5.1 (Friard & Gamba, 2016). Relevant behaviours were recorded as either “point events” or “states” (to quantify the number or duration of events, respectively; see ethogram in Table 1) and the identity and sex of each individual was determined from its unique colour ring combination. In a minority of cases, rings were not visible in the video during a bird’s visit to the nest box, so the individual’s sex was recorded as “unknown”. When the sex was relevant for analyses, we excluded data from unknown focal individuals. If vocalisations occurred when both members of a pair were in the nest box, we used fine-scale body movement (e.g. of beak or thorax) to establish which individual was vocalising. We analysed different types of vocalisations with distinct acoustic qualities separately because they may be functionally different. We analysed “chatter”, a distinctive sequence of repeated high pitch vocalisations, separately from other calls (hereafter called “calls”) as they may be functionally different.

**Table 1.**
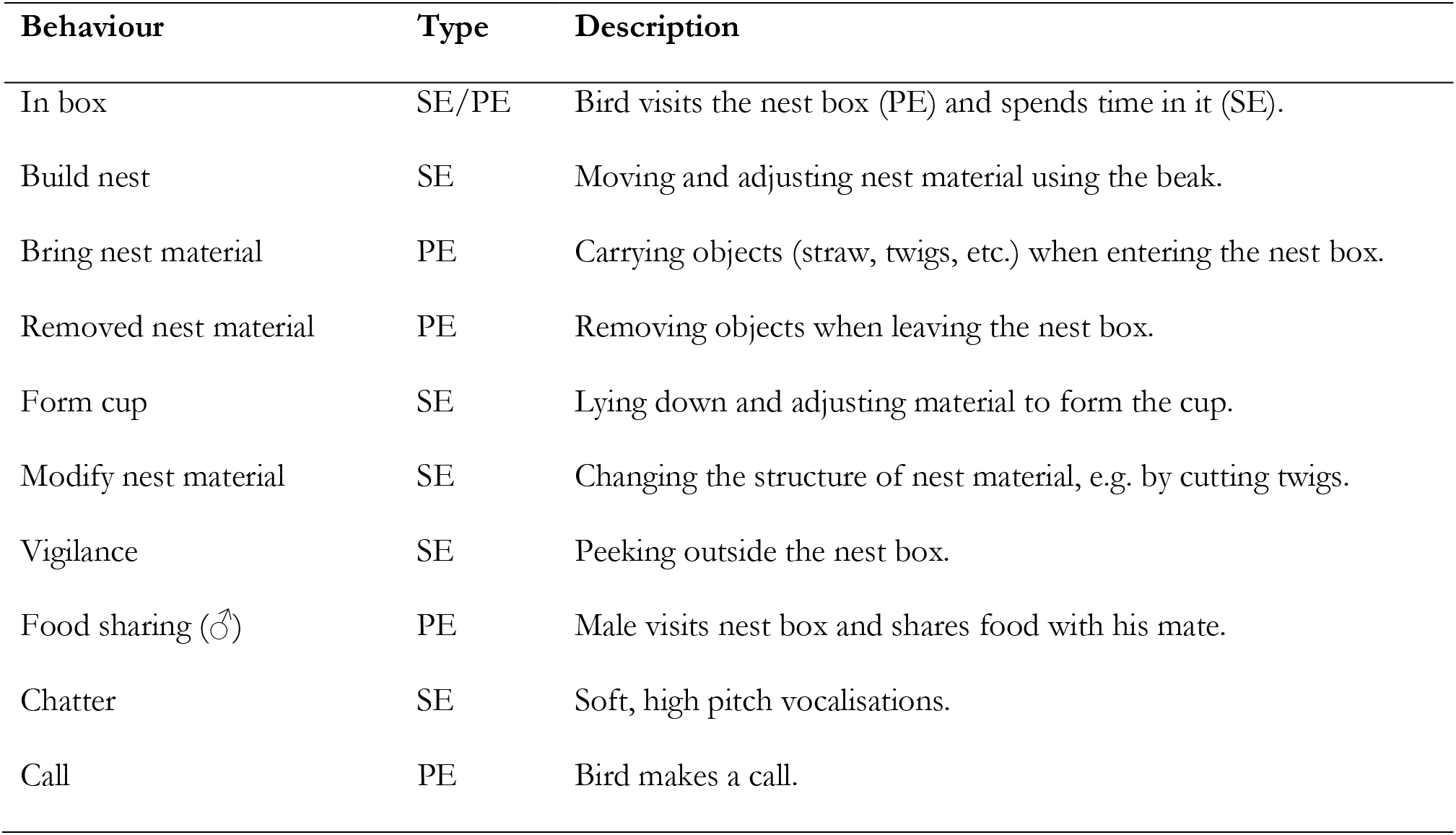
Ethogram of behaviours recorded. Type denotes whether a behaviour was a point event (PE) or a state event (SE) with a duration. Begging calls by females are included in the event “call”.

### Statistical analyses

Data were analysed in *R* version 4.0.2 (R Core Team, 2020). For all behavioural variables (N = 10), we extracted raw total durations (states) and quantities (point events) per observation for each pair (in total) and for both sexes separately. To standardise variables, we divided raw data (durations or counts) by the observation length (in seconds), and multiplied state events by 100 (percentage of time) and point events by 3600 (rate per hour). For each standardised behavioural variable, we calculated means and standard deviations (denoted as mean ± SD henceforth) across pairs.

### General procedure: mixed models and repeatability analysis

#### Mixed models

We analysed data with (generalised) linear mixed models (LMMs and GLMMs), using the packages *lme4* (for LMMs) (Bates et al., 2015) and *glmmTMB* (for GLMMs) (Brooks et al., 2017). To formulate models, we selected the dependent variable of interest (a behaviour or a fitness proxy) and one or more biologically meaningful independent variable(s). We subsequently tested model assumptions such as normality of residuals, homoscedasticity, error structure, dispersion, zero inflation, and influential datapoints (Cook’s distance), using diagnostic plots and tests implemented in *R* (LMMs) or in the package *DHARMa* (GLMMs) (Hartig, 2019). To infer estimates and P-values, we used Wald tests in the package *car* (Fox & Weisberg, 2011). All models included the variables ‘year’ (fixed effect), ‘pair ID’ (random effect), and ‘study site’ (random effect) to account for temporal and spatial variation as well as pseudoreplication. We had no specific a priori predictions as to the effects of the birds’ age, but as it could potentially influence behaviour and reproductive success, we initially included ‘age’ (in years) in analyses as an additional fixed effect. If age did not appear to play an important role, we removed the variable from analyses to avoid over-parametrisation and maximise statistical power. Observation-level random effects (X. A. Harrison, 2014) and COM-Poisson error structures accounted for zero-inflation and underdispersion, respectively.

#### Repeatability analysis

We calculated the repeatability of behaviours and fitness proxies in pairs for which repeated measures were available (N = 16 pairs observed more than once within and/or across years), using the package *rpt*R (Stoffel et al., 2017). When quantifying repeatability of state events, we used Box-Cox transformations (Sakia, 1992) to meet assumptions of Gaussian data.

### Sex differences in behaviour

We could identify birds for 76.71 ± 30.48 percent of the time spent in the box. To quantify sex differences in behaviour, we used a subset of behavioural data where the identity of the focal individual(s) was known (N = 62 videos of 40 pairs; two cases were removed in analyses including vocal communication because the microphones failed to record: box Y02, 2018 and box Z19, 2018). We investigated the time both sexes invested in ‘vigilance’, ‘nest building’, ‘being in the nest box’, ‘forming the cup’, and ‘chatter’, using separate LMMs, with the standardised response variables log-transformed to meet model assumptions (West et al., 2014). We also examined whether either sex invested more time in ‘vigilance’ or ‘nest building’ as response variables given that these were the most frequent behaviours in the nest box. Models examining only vigilance as response terms also contained the ‘number of days the video had been recorded before the lay date’ (covariate) to account for the potential influence the date may have on behaviours. For instance, birds may reduce their effort closer to the lay date when the nest should be completed. On the other hand, males could increase their vigilance closer to the lay date to guard the female during her fertile period. ‘Modification of material’ was too rare to permit formal statistical analysis. We also conducted separate GLMMs on rates of ‘material brought’, ‘material removed’, ‘visits to the nest box’, and ‘calls’ (rounded to rates per hour and treated as count data) fitted as a response term. In these analyses, ‘sex’ was the main predictor variable of interest, but we also modelled an interaction between sex and age of each bird to examine whether sex differences may be age-dependent and also to include age as a covariate potentially affecting behaviour.

### Nest building and reproductive success

#### Dependent variables: reproductive parameters

To examine fitness correlates of behaviours, we separately analysed three different proxies for reproductive success (Table 2). Firstly, we used the relative lay date of the first egg compared to the date the first clutch was initiated per site. As colonial breeders, jackdaws breed within a relatively short time period, and early layers may benefit from lower competition with other colony members. A second proxy of reproductive success was the clutch size. Thirdly, we examined the volume of the first and the third egg). Jackdaws lay an egg per day until the clutch is complete, and they show hatching asynchrony, with the first egg being the one which is most likely to survive. The second egg has a relatively high probability to survive as well, whereas the survival rate of the third egg is approximately 0.5. We did not analyse later eggs as these rarely survive (Goumas et al., In prep.). One pair (box Z28, 2019) was excluded from these analyses because it was displaced by another pair during nest construction so could not produce a clutch. When analysing egg volume, we removed one pair (box Y21, 2019), which had been parasitised by a conspecific female. Two pairs were removed when analysing the third egg volume as they only laid two eggs (box Y16, 2019; box Z15, 2019).

**Table 2.** Reproductive parameters to study fitness correlates of behaviour. “N” denotes the sample size (number of videos and of pairs, respectively) used in the models (unless stated otherwise). To calculate mean, standard deviation, and the range of parameters, the sample size was smaller compared to the numbers presented in this table because seven pairs observed twice in one year were only considered once here.

| Reproductive parameter | Definition | Error structure | N | Mean ± SD | Range (min. – max.) |
| --- | --- | --- | --- | --- | --- |
| Lay date | Day first egg laid (1 = first egg per site) | COM-Poisson | 61 (39) | 5.19 ± 2.92 | 1 - 14 |
| Clutch size | Number of eggs laid in total | COM-Poisson | 61 (39) | 4.43 ± 0.87 | 2 - 6 |
| Egg volume | Egg volume of first egg (cm<sup>3</sup>) | Gaussian | 60 (38) | 11.42 ± 0.85 | 9.75 - 13.19 |
| Egg volume | Egg volume of third egg (cm<sup>3</sup>) | Gaussian | 58 (36) | 11.23 ± 1.08 | 8.62 - 13.35 |

#### Behavioural predictors

We defined four ‘behavioural concepts’ to be used separately as independent variables that may relate to measures of reproductive success. For each of the first three concepts we calculated a distinct PCA to summarise (scaled) behavioural variables to be included in models whilst minimising model complexity (Budaev, 2010; Morton & Altschul, 2019) and to account for multicollinearity among variables (Graham, 2003). When performing a PCA, we calculated a correlation matrix including the variables of interest, applied the KMO-measure (threshold 0.5) to test for sufficient correlation among them (Budaev, 2010), and conducted a parallel analysis to determine the number of principal components to be considered objectively (Morton & Altschul, 2019), using the package *psych* (Revelle, 2018). We constructed alternative models in cases where we analysed distinct predictors each of which reflected a specific hypotheses (for more details see below). To select a model, we employed Akaike’s Information Criterion (*AICc* to account for small sample sizes (X. A. Harrison et al., 2018)) in the package *bbmle* (Bolker & R Core Team, 2017). A model with the lowest *AICc* had to differ at least 2 *AICc* units to be selected. In instances where only one predictor variable corresponded to one of the behavioural concepts, we did not use model selection and constructed a single model per fitness variable instead. When we detected a significant relationship between behavioural predictors and a fitness proxy, we performed models again with a subset of observations for which data on female body condition was available in that particular year to control for this variable (covariate). We then compared two models with and without female body condition using likelihood ratio tests. Body condition was quantified using the residuals of a regression examining the relationship between measures of body size (tarsus and wing length) as independent and body weight as dependent variables. In all models investigating fitness correlates of behaviour, we also included the ‘number of days the video was recorded before the lay date’ (‘day’ henceforth, covariate), because this could have influenced the birds’ behaviour. Moreover, we fitted ‘female age’ (covariate) to account for breeding experience (female and male age was significantly correlated: *ϱ* = 0.744, t_59_ = 8.560, *P* < 0.001). Another covariate was ‘food sharing’ by males, because this cooperative behaviour not directly related to nest building could affect reproductive success. We outline each concept, and the analytical methods used to examine it, below.

##### (i) Overall activity levels and intensity of behaviours

To test prediction 3 that pairs investing more in the nest relative to other activities, such as vigilance, should lay earlier, and have larger clutches and eggs, we constructed a PCA of nine behavioural variables (‘PCAAll’)(Table 3; Table A 4). The variable ‘food sharing’ (rate per hour) was left out in the PCA due to the KMO-threshold (0.43), but included in the models as a covariate as levels of food sharing by the male could influence the female’s ability to invest in the nest and the clutch. All behavioural variables loaded negatively onto PC1_All_, which could therefore reflect ‘intensity’ of behaviours. The four nest building behaviours loaded negatively onto PC2_All_, whereas the other behaviours (vigilance, vocalisations, time in the box) loaded positively onto PC2_All_. These opposite loadings suggest a trade-off, such that pairs may have invested relatively more time in either the nest or in vigilance and vocalising on the other. Therefore, we hypothesised that (i) all behaviours (PC1_ALL_), (ii) a relative investment in nest building compared to other behaviours (PC2_All_), or (iii) both (PC1_ALL_ and PC2_ALL_) could be used as predictor of reproductive success. We formulated three corresponding models and two further models which contained (iv) only ‘year’ and ‘day’ and (v) only an ‘intercept’. Subsequently, we compared these models using AICc.

**Table 3.**
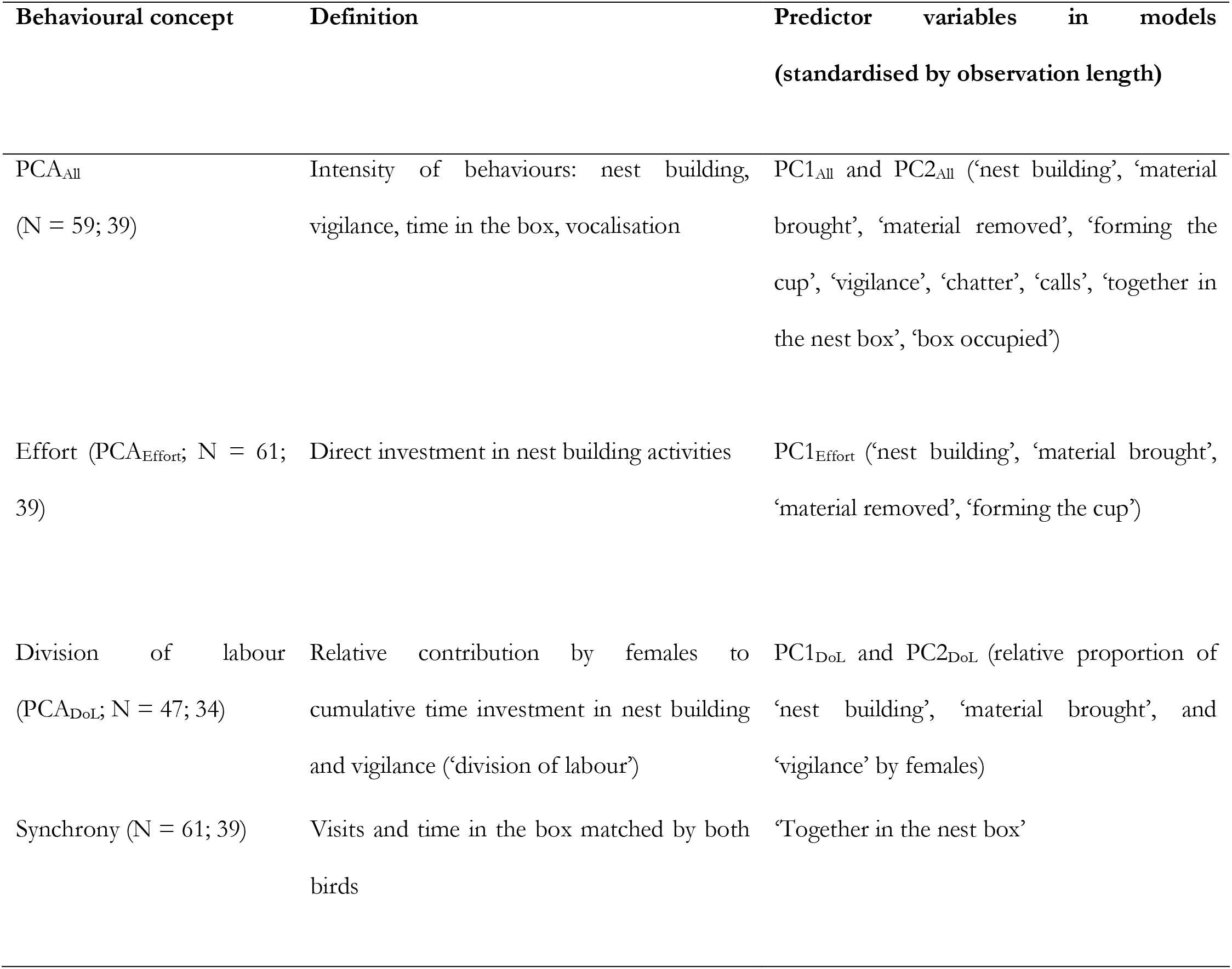
Behavioural concepts used to examine correlates of reproductive success. The two sample sizes denote the number of observations and the number of pairs, respectively. In the PCA_All_, the variables ‘food sharing’ (KMO = 0.46) and ‘modify’ (rare behaviour) were left out. PC2_DoL_ was equivalent to ‘material brought’ by females (loading 0.99) and therefore the latter variable was used. For more details on the PCA please refer to the Appendix.

##### (ii) Direct investment in the nest (‘effort’)

To examine prediction 4, we analysed the relationship between a PCA comprising variables directly related to nest building and reproductive success (‘PC1_Effort_’; Table 3; Table A 5). All four variables loaded negatively onto ‘PC1_Effort_’, suggesting it could be used as a measure of total nest building effort. Following the parallel analysis that quantified the number of principal components to be used, we constructed only one model per fitness proxy, including ‘PC1_Effort_’ as an independent variable instead of comparing alternative models.

##### (iii) Relative investment by females in nest building and vigilance (‘division of labour’, ‘DoL’)

We conducted a third PCA to examine ‘division of labour’ (‘DoL’) (prediction 5), that is, whether the relative proportion of female contribution to nest building and vigilance (compared to the sum of female and male effort) was linked to reproductive success (‘PCA_DoL_’; Table 3; Table A 6). In this analysis, the sample size was smaller (N = 47 observations), because we discarded observations when a proportion could not be calculated (neither sex of a pair showed one of the behaviours). PC1_DoL_ suggested that females contributed either more through nest building or vigilance due to opposite loadings (Table A 6). PC2_DoL_ was equivalent to the ‘relative proportion of material brought by females’ (loading = 0.99; Table A 6). When analysing ‘PCA_DoL_’, we constructed alternative models per fitness proxy by using either (i) ‘PC1_DoL_’, (ii) the ‘relative proportion of material brought by females’, (iii) only ‘year’ and ‘day’, or (iv) only an ‘intercept’ as independent variables. For predictors (i) and (ii) we also modelled a quadratic effect which could indicate that equal contributions by both sexes are related to greater reproductive success.

##### (iv) Time spent together in the nest box (‘synchrony’)

To test prediction 6 that the level of ‘synchrony’ could be linked to reproductive success, we used the ‘proportion of time individuals spent together in the nest box’ as an independent variable. To examine its relationship with fitness measures, we constructed one model per fitness proxy with ‘synchrony’ being the only independent variable.

## 3. Results

### Behaviours and sex differences

#### Sex differences

On average, jackdaw pairs occupied their nest box for 29.09 ± 19.64 percent of the observations and spent 23.17 ± 24.58 percent of that time together. The behavioural repertoire of both sexes was broadly similar (Figure 1), but they also differed in some behaviours (Figure 2, Table 4). Specifically, females spent 1.5 times more time building the nest than males (Figure 2a, Table 4). Males spent on average 1.4 times more time being vigilant than females, but this difference was not significant (Figure 2b, Table 4). Moreover, males spent more time being vigilant than they spent building, whilst females were similar in both behaviours (Table 4). Males did not increase vigilance when the observation was recorded closer to the lay date (LMM, sex * days before lay date: β ± SE = 0.021 ± 0.023, *X*^2^_1_= 0.887, 95 % CI [− 0.01, 0.03], *P* = 0.346). Females called 1.9 times more frequently than males (Figure 2c, Table 4), even after removing female begging calls (GLMM, sex: β ± SE = – 0.690 ± 0.228, *X*^2^_1_= 9.120, *P* = 0.003). There was weak evidence that older birds brought more nesting material (GLMM, sex: sex: β ± SE = 0.257 ± 0.124, *X*^2^_1_= 4.135, 95 % CI [0.013, 0.500], *P* = 0.042; Figure A 1), but this relationship was not maintained when removing the four youngest individuals that were two years old (GLMM, age: β ± SE = 0.177 ± 0.127, *X*^2^_1_= 1.645, 95 % CI [− 0.073, 0.426], *P* = 0.200). Aside from this there was no evidence for any age effects or sex by age interactions on any aspect of building behaviour (Table A 3).

**Figure 1.**
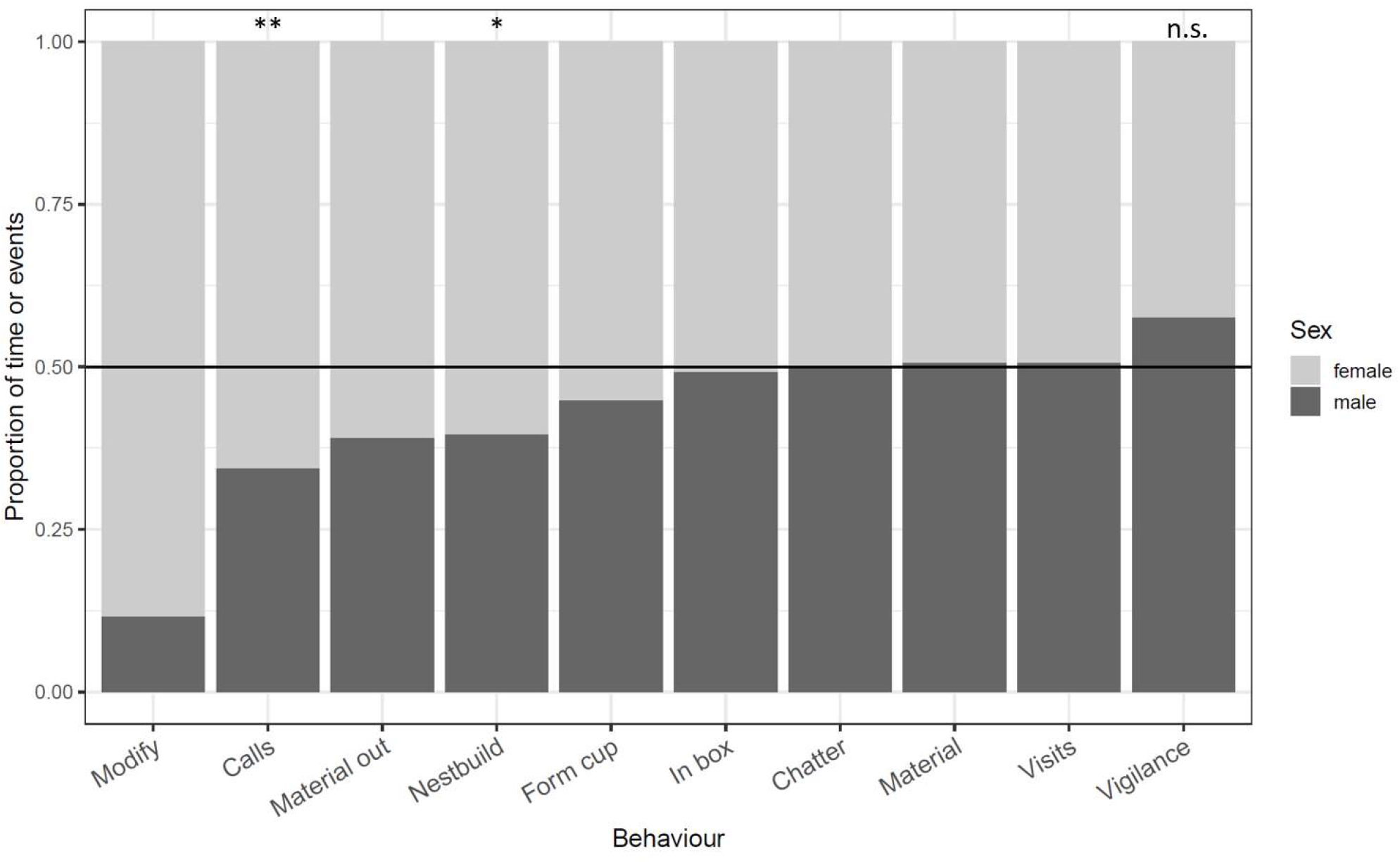
Mean relative duration (state events) and frequency of events (point events) by sex (N = 62 observations; N = 60 for vocalisations). The horizontal line marks the proportion of 0.5, meaning both sexes showed a behaviour equally long or often, respectively. Asterisks indicate a significant sex difference in behaviour based on the model output (calls, nest building; < 0.01 ** and < 0.05 *).

**Figure 2.**
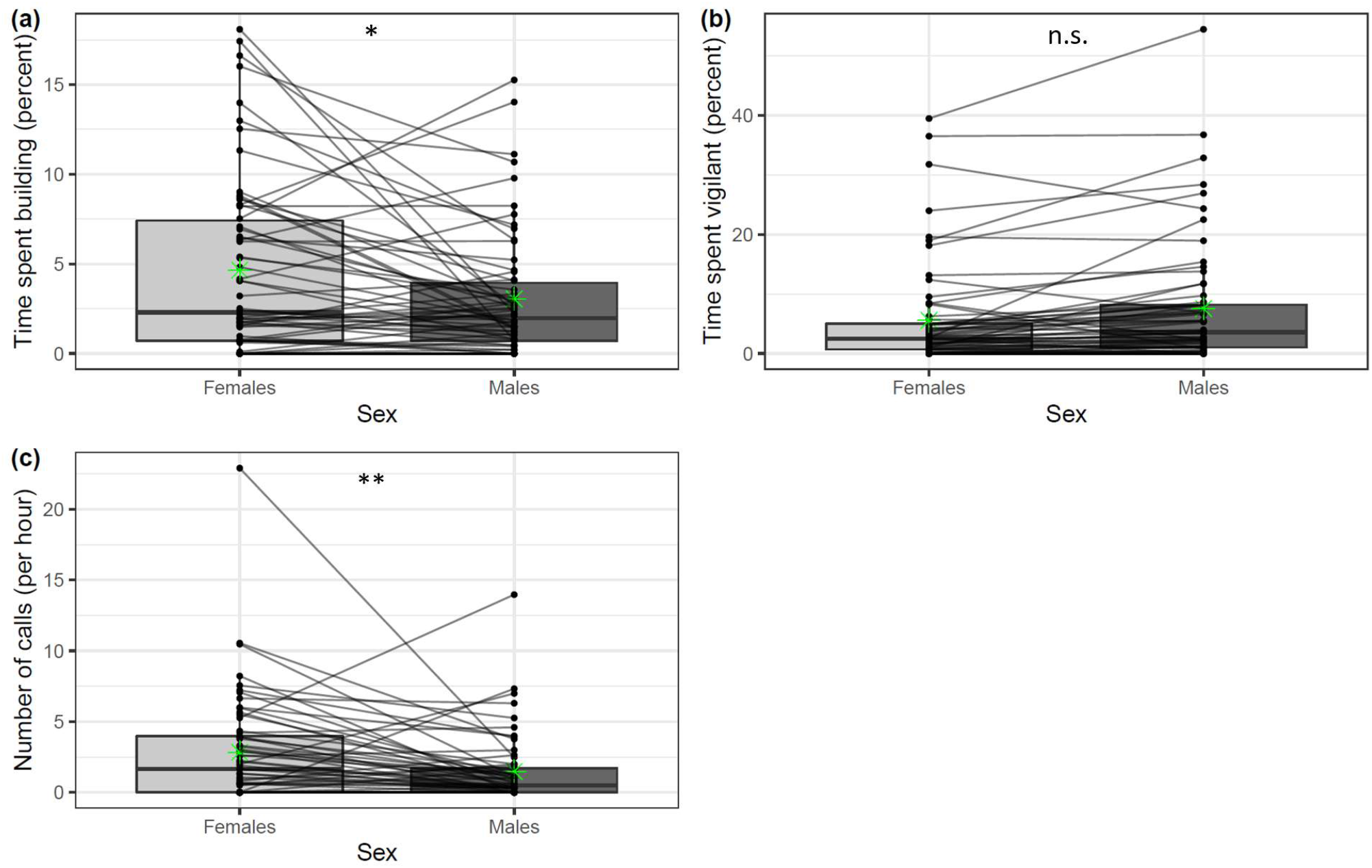
Sex differences in three behavioural variables (N = 62 observations; N = 60 for calls). Horizontal lines in the boxes indicate the median, green asterisks indicate the mean. Upper and lower ends of the boxes reflect the 0.25 and the 0.75 quartiles, respectively. Horizontal lines connecting points represent distinct pairs. Asterisks indicate a significant difference (** < 0.01; * < 0.05; n.s. = 0.060). **(a)** The time spent building the nest as a percentage of the observation length. **(b)** The time spent being vigilant as a percentage of the observation length. **(c)** The number of calls made by both sexes (per hour).

**Table 4.**
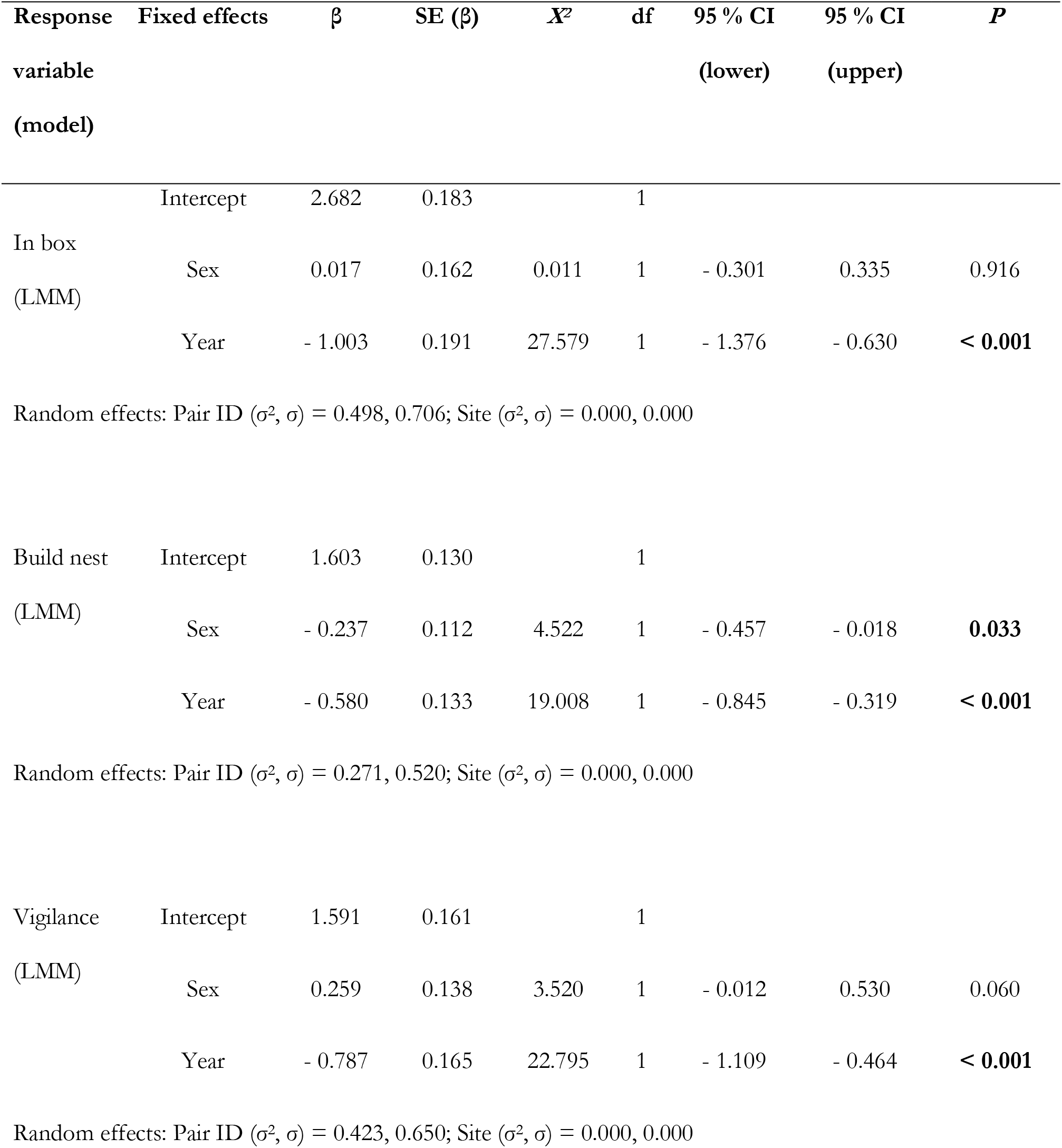
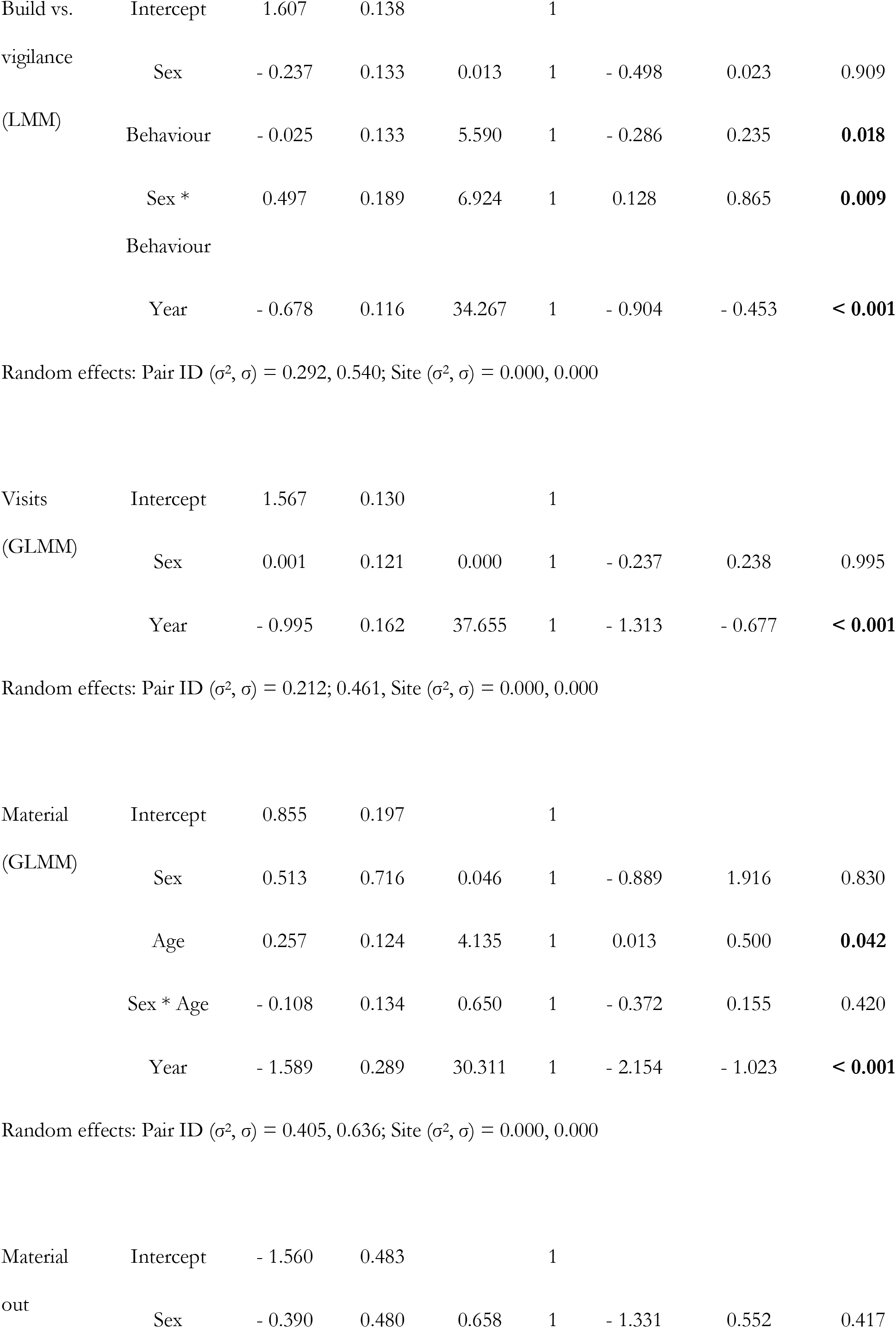
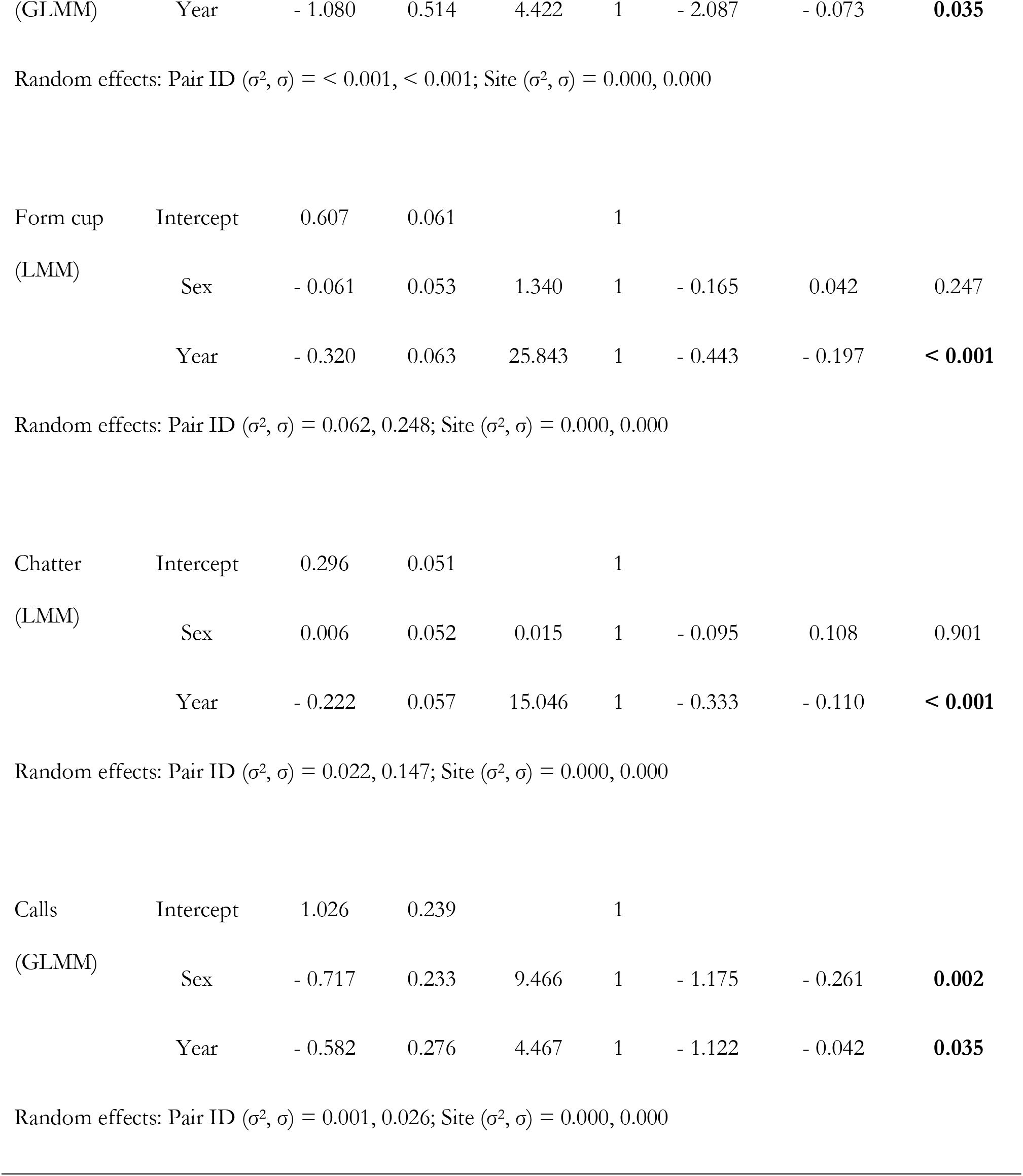
Sex differences in behaviour. Statistically significant results are in bold. Response variables for LMMs were log-transformed and estimates for GLMMs (COM-Poisson) are on the log scale. Intercepts refer to the values of females and to the year 2018. Observation-level random effects accounted for zero-inflation. σ^2^ and σ denote the variation and standard deviation attributed to random effects. Sex differences were not examined for rarer behaviours (modification of nest material).

#### Repeatability and variation across pairs and years

There was considerable variation in behaviours between different pairs (Figure 2; Table A 1; Table A 2), but when inside the nest box, the majority of their time was spent building the nest or being vigilant (Table A 1). On the level of the pair, birds that spent more of their time in the nest box together spent more time being vigilant (*ϱ* = 0.906, t_60_ = 16.547, *P* < 0.001), but not more time building (*ϱ* = 0.139, t_60_= 1.090, *P* = 0.280). Conversely, pairs in which only one individual occupied the nest box for longer spent more time building (ϱ = 0.840, t_60_ = 11.996, *P* < 0.001), but not more time being vigilant (ϱ = 0.175, t_60_ = 1.379, *P* = 0.173). The time females spent building and males spent being vigilant was positively correlated (ϱ = 0.271, t_60_ = 2.184, *P* = 0.033). No behaviour was repeatable in 16 pairs for which repeated measures were available within and/or between years (Table A 8). Jackdaws varied in their behaviour depending on the year (Table 4).

#### Behaviours and correlates of reproductive success

The volume of the first egg (R = 0.598 ± 0.176, 95 % CI [0.182, – 0.850], *P* = 0.009) and the third egg (R = 0.598 ± 0.187, 95 % CI [0.115, – 0.848], *P* = 0.007) females laid was repeatable across years. The relative lay date of pairs compared to the first lay date per site was also repeatable (R = 0.643 ± 0.204, 95 % CI [0.018, – 0.822], *P* = 0.015). Clutch size was not repeatable for those pairs observed in both years (Table A 9).

#### Lay date

The majority of jackdaw females laid their first egg in the middle of April (17.05 ± 3.20 days where 1 = 1^st^ April; 5.19 ± 2.92 days after the first clutch was initiated per site). Females that laid their first egg relatively earlier contributed relatively more to bringing material (Figure 3; Table 5). In this model, female age did not appear to have an effect and was therefore removed (β ± SE = 0.086 ± 0.082, *X*^2^_1_= 1.108, *P* = 0.292, 95 % CI [− 0.074, 0.246]). Including a proxy for female body condition did not improve the model (*X*^2^_1_ < 0.001, *P* = 1.0).

**Figure 3.**
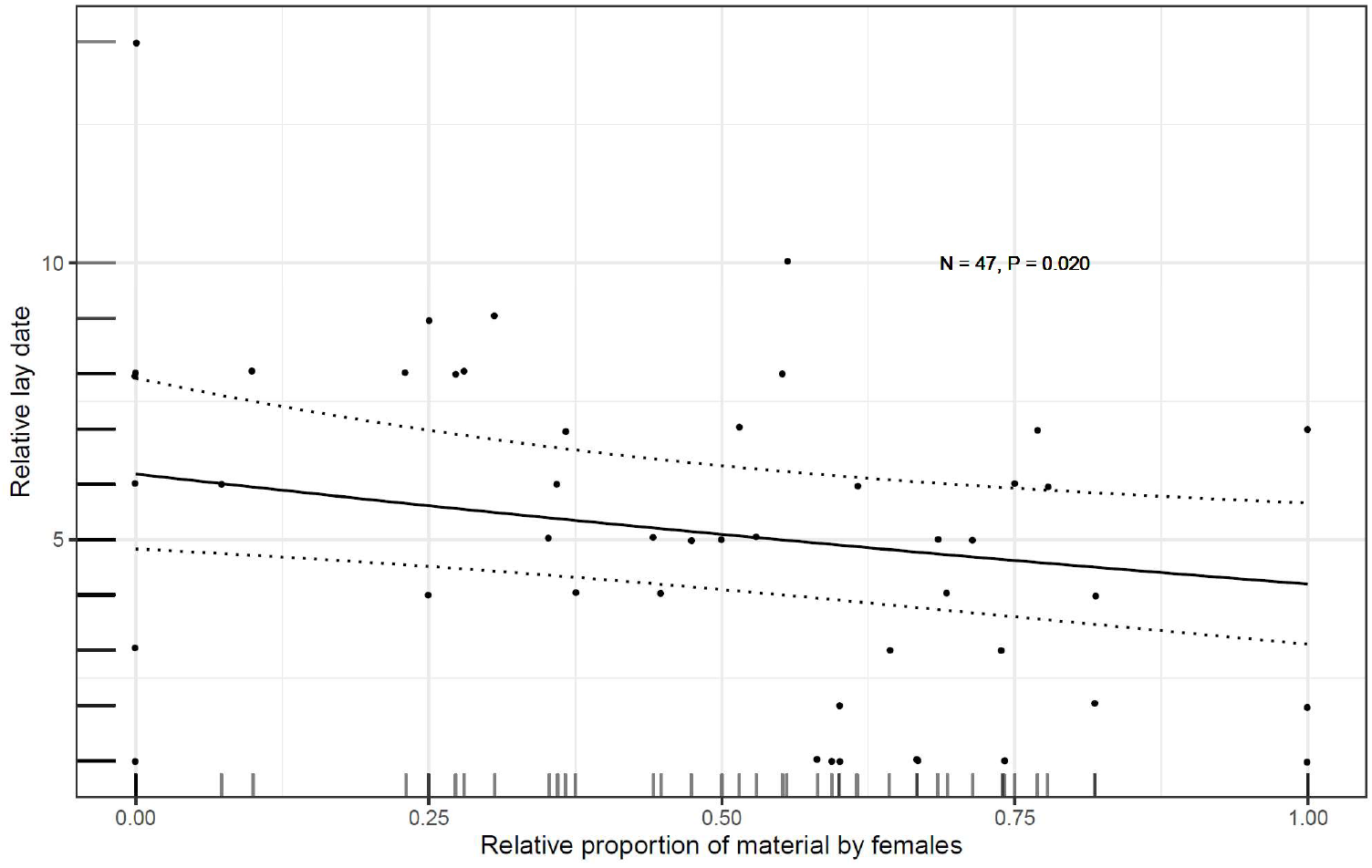
Relationship between the relative lay date (number of days compared to the day the first clutch was initiated per site) and female contribution to transporting nest material to the nest box. The relative proportion of items brought by females refers to the total amount of nest material brought by females and males. Dots indicate raw data; dotted lines show the 95 % confidence intervals around the fitted line (solid) from the model output.

**Table 5.**
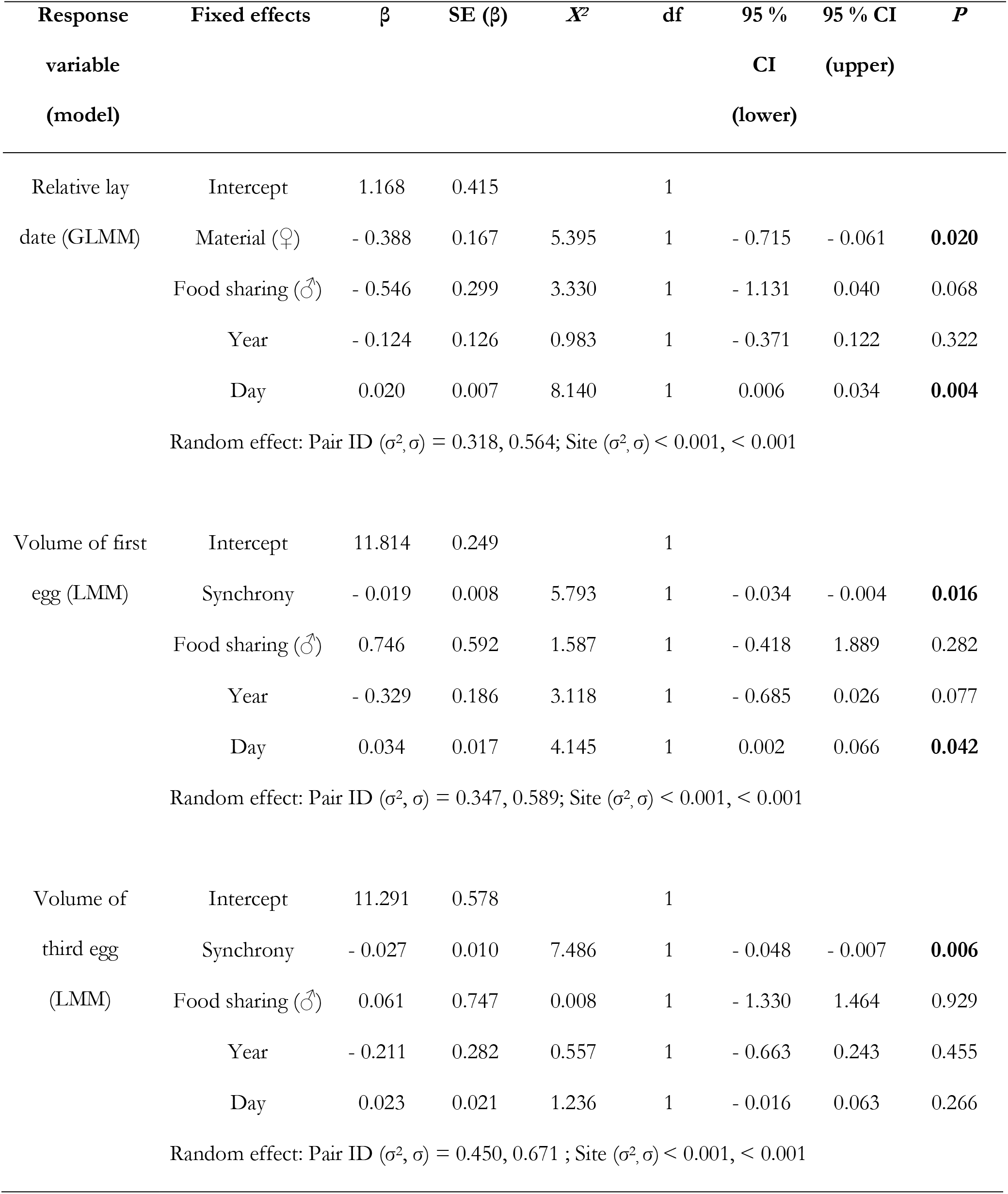
Summary of the statistical analyses on the relationship between behaviours at the nest building phase and fitness proxies. The fixed effect ‘Material (♀)’ reflects the relative female contribution to bringing nest material relative to the overall effort by both sexes. The reference year was 2018 and ‘day’ refers to the number of days the video was recorded before the lay date of the first egg. σ^2^ and σ show the variation and standard deviation explained by random effects. This table shows significant results; further details are in the Appendix (Table A 7).

| Response | Fixed effects | $\beta$ | SE ($\beta$) | $X^2$ | df | 95 % | 95 % CI | $P$ |
| --- | --- | --- | --- | --- | --- | --- | --- | --- |
| variable |  |  |  |  |  | CI | (upper) |  |
| (model) |  |  |  |  |  | (lower) |  |  |
| Relative lay | Intercept | 1.168 | 0.415 |  | 1 |  |  |  |
| date (GLMM) | Material (♀) | - 0.388 | 0.167 | 5.395 | 1 | - 0.715 | - 0.061 | **0.020** |
|  | Food sharing (♂) | - 0.546 | 0.299 | 3.330 | 1 | - 1.131 | 0.040 | 0.068 |
|  | Year | - 0.124 | 0.126 | 0.983 | 1 | - 0.371 | 0.122 | 0.322 |
|  | Day | 0.020 | 0.007 | 8.140 | 1 | 0.006 | 0.034 | **0.004** |
| Random effect: Pair ID ($\sigma^2, \sigma$) = 0.318, 0.564; Site ($\sigma^2, \sigma$) < 0.001, < 0.001 | | | | | | | | |
| Volume of first | Intercept | 11.814 | 0.249 |  | 1 |  |  |  |
| egg (LMM) | Synchrony | - 0.019 | 0.008 | 5.793 | 1 | - 0.034 | - 0.004 | **0.016** |
|  | Food sharing (♂) | 0.746 | 0.592 | 1.587 | 1 | - 0.418 | 1.889 | 0.282 |
|  | Year | - 0.329 | 0.186 | 3.118 | 1 | - 0.685 | 0.026 | 0.077 |
|  | Day | 0.034 | 0.017 | 4.145 | 1 | 0.002 | 0.066 | **0.042** |
| Random effect: Pair ID ($\sigma^2, \sigma$) = 0.347, 0.589; Site ($\sigma^2, \sigma$) < 0.001, < 0.001 | | | | | | | | |
| Volume of | Intercept | 11.291 | 0.578 |  | 1 |  |  |  |
| third egg<br>(LMM) | Synchrony | - 0.027 | 0.010 | 7.486 | 1 | - 0.048 | - 0.007 | **0.006** |
|  | Food sharing (♂) | 0.061 | 0.747 | 0.008 | 1 | - 1.330 | 1.464 | 0.929 |
|  | Year | - 0.211 | 0.282 | 0.557 | 1 | - 0.663 | 0.243 | 0.455 |
|  | Day | 0.023 | 0.021 | 1.236 | 1 | - 0.016 | 0.063 | 0.266 |
| Random effect: Pair ID ($\sigma^2, \sigma$) = 0.450, 0.671 ; Site ($\sigma^2, \sigma$) < 0.001, < 0.001 | | | | | | | | |

#### Clutch size

Females laid a mean of 4.43 ± 0.87 eggs. Clutch size was unrelated to any of the behavioural predictors (Table A 7).

#### Egg volume

The mean volume of the first and the third egg was 11.42 ± 0.85 cm^3^ and 11.23 ± 1.07 cm^3^, respectively. The volume of both the first and the third egg was smaller in birds that spent more time together in the box (Figure 4; Table 5). This relationship remained after excluding an influential datapoint (a pair that spent more than 60 % of the time together in the nest box) (LMM, synchrony, first egg: β ± SE = – 0.027 ± 0.012, *X*^2^_1_= 5.590, *P* = 0.018, 95 % CI [− 0.051, − 0.003]; third egg: β ± SE = − 0.046 ± 0.014, *X*^2^_1_= 10.648, *P* = 0.001, 95 % CI [− 0.075, – 0.016]). In the models examining synchrony there was no effect of female age on egg volume (LMM, age, first egg: β ± SE = 0.102 ± 0.094, *X*^2^_1_= 1.169, *P* = 0.280, 95 % CI [− 0.076, 0.290]; third egg: β ± SE = 0.005 ± 0.115, *X*^2^_1_= 0.002, *P* = 0.962, 95 % CI [− 0.212, 0.228]. Including female body condition did not improve the model examining the relationship between synchrony and first egg volume (*X*^2^_1_ 0.113, *P =* 0.945), and between synchrony and third egg volume (*X*^2^_1_ 0.013, *P* = 0.909).

**Figure 4.**
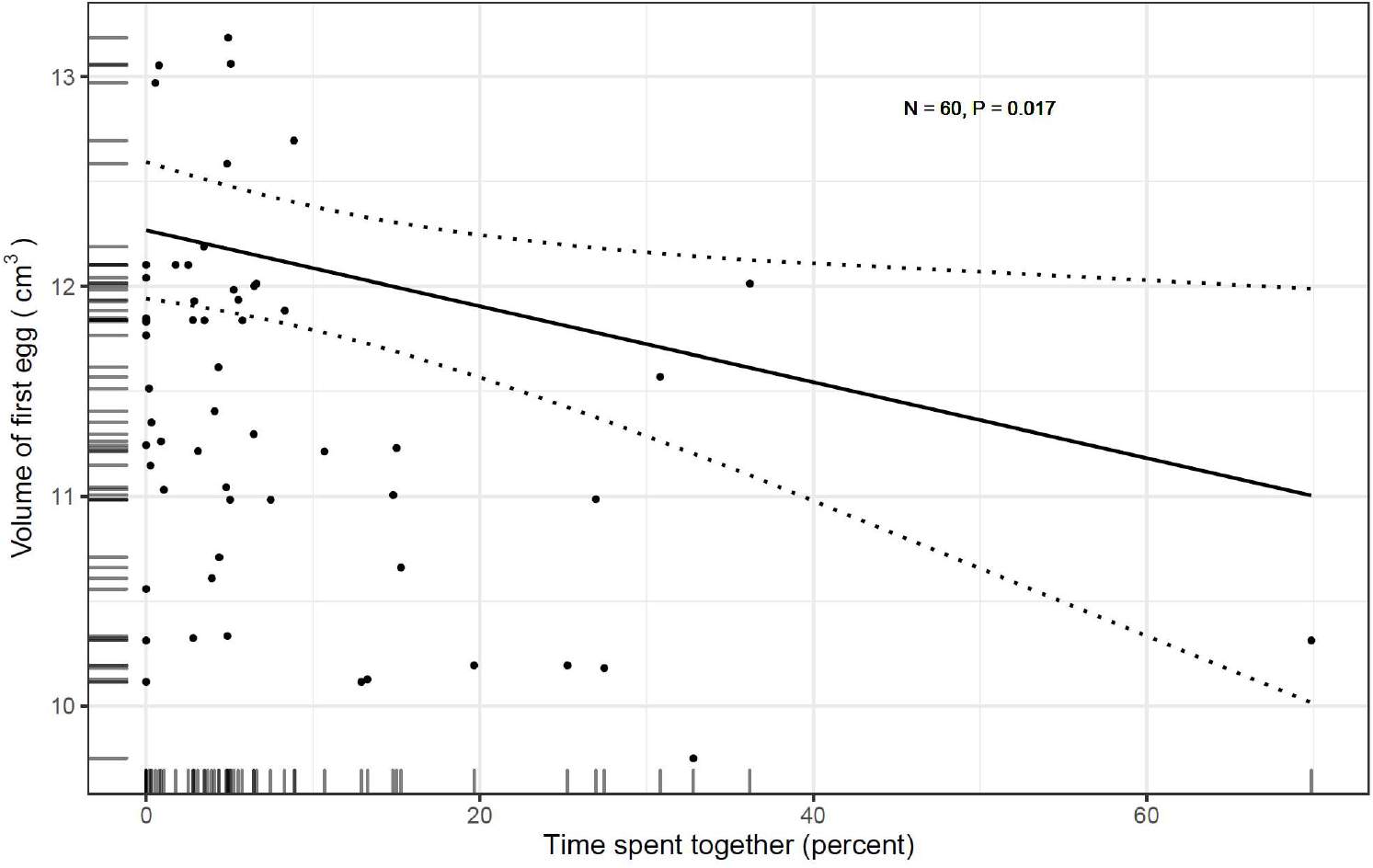
Relationship between the first egg volume per female and the percentage of time mates spent together in the nest box. The continuous fitted line corresponds to the model output; dotted lines show the 95 % confidence intervals. The result was still significant when excluding an outlier (a pair spending more than 60 % of the time together in the box).

## Discussion

Our findings demonstrate that in monogamous jackdaws nest building entails substantial investment from both partners, and may have important fitness consequences. Although both sexes exhibited a similar behavioural repertoire and cooperated to create their nest, there were some sex differences in investment, with females building more and males tending to be more vigilant. Moreover, some aspects of behaviour, such as the relative female contribution to transporting material and time spent together in the box, were associated with measures of reproductive success (lay date and egg volume).

Consistent with our predictions, jackdaws cooperated during nest construction, with the two sexes behaving broadly similarly, with both investing in bringing material, building the nest, and forming the cup. In jackdaw pairs fitness outcomes are interdependent due to repeated mating opportunities with the same partner across years and low rates of divorce and successful extra-pair copulations (Gill et al., 2020; Wechsler, 1989). Under these circumstances, conflicts of interest between mates may be minimised, particularly if biparental care is necessary to successfully rear offspring, ultimately favouring cooperation. Although nest building by jackdaws clearly requires a substantial cooperative investment from both partners, our findings suggest moderate division of labour could facilitate cooperation (cf. Iserbyt, Fresneau, Kortenhoff, Eens, & Müller, 2017). Females built more than males and were therefore more responsible for the nest structure. In contrast, males dedicated more time to vigilance than to building, which may be particularly important in colonially nesting jackdaws, where intraspecific competition over nest cavities is severe and can constrain reproduction (Henderson & Hart, 1993; Röell, 1978; Verhulst & Salomons, 2004). Vigilant residents may not only anticipate threatening non-resident competitors searching for a nest cavity, but their bright eye colour has also been shown to deter intruders (Davidson et al., 2014). Males may prioritise vigilance because the risks of vigilance and defence may be more costly for females as they need to stay in a good condition for later stages of breeding, such as incubation. Additionally, males may invest relatively more in vigilance than in building due to their slightly larger body size (Fletcher & Foster, 2010), a trait which impacts contests in this species (Verhulst et al., 2014). There was no significant difference in the amount of time males and females spent in vigilance, and male vigilance was independent of days until his partner’s fertile window (beginning 5 days prior to the lay date; Gill et al., 2020), suggesting that vigilance serves primarily to defend the nest site rather than as a form of mate-guarding. Males cooperated, for example through vigilance and transporting nest material despite contributing less to building the nest by arranging material in the nest box. By increasing their own nest building activity, females may be able to partially compensate for this. Females may also spend more time building than males because they may be better informed about their own requirements for incubating the clutch. The mechanisms through which partners acquire and act upon information to respond to each other’s behaviour and coordinate division of labour remains unknown. Elucidating these mechanisms will be vital to understanding the cognitive demands of pair-bonding, such as the need to track and respond to another’s behaviour (Emery et al., 2007).

Our results suggest substantial variation in behaviour and time budgets between pairs. Furthermore, no behavioural variable was significantly repeatable within pairs, indicating there may also be considerable behavioural variation within pairs. It is possible that the lack of repeatability within pairs is an artefact of differences in sampling between years, because videos were recorded significantly closer to the lay date in 2019, which could have affected the behaviour. For instance, pairs may have seemingly built less in 2019, but this could have been because the video was recorded closer to the lay date. Given the limited amount of data per pair and the fact that not all pairs were observed repeatedly, these results must be interpreted with caution. Nevertheless, our findings raise the possibility that there may be substantial phenotypic plasticity in jackdaw nest building behaviour, in keeping with recent evidence nest building behaviour may be less ‘fixed’ than previously thought (Walsh et al., 2013). Indeed, we found intensity of behaviours significantly varied across two years, implying that environmental variables may affect behaviour, and also measures of reproductive success.

Some behaviours during nest construction were associated with proxies for reproductive success, raising the possibility that selection pressures may act on how pairs cooperate and how they spend their time. The relative contribution of females to bringing nest material was associated with an earlier lay date. Given that early laying can reduce competition for food when provisioning offspring and is often linked to elevated reproductive success in birds (Perrins, 1970), this suggests the female contribution to nest building may have important fitness consequences. We had hypothesised that more equal contributions by both partners could enable an earlier lay date by reducing the time and energy needed to build the nest, potentially important for females to save energy for costly egg production (Williams, 2005). Instead, we found that the time females spent building and males spent being vigilant was positively correlated, suggesting that greater investment in vigilance by the male may allow the female to invest more energy in building the nest and thus lay earlier.

The amount of time partners spent together was also linked to fitness outcomes, but in the opposite direction to our predictions. Whereas we had predicted that greater synchrony would reflect compatibility between partners and be linked to reproductive benefits (Spoon et al., 2006), we actually found more synchronous pairs laid smaller eggs. One possible explanation for this is that the pairs that spent more time together in the nest were those that faced greater competition, as both partners are required to successfully guard a nest site in this species (Röell, 1978; Verhulst & Salomons, 2004). Indeed, we found that pairs that spent more time together invested more time in vigilance but not in building the nest. This suggests a competitive and stressful period where the need to defend the nest box detracts from investment in nest building (Röell, 1978). There is evidence from other species, such as house sparrows (*Passer domesticus),* that investment in parental care, and consequent reproductive success, can be impaired by chronically elevated stress hormone levels (Ouyang et al., 2011). While including morphological measures of female body condition did not improve our models, measures of current energetic and physiological state may prove more useful in future studies.

Together, our findings indicate that nest building in monogamous birds provides an important, but as yet understudied, model system to investigate the evolution and proximate mechanisms of cooperation. How much a partner invests in nest building may be a source of information used by individual birds to assess how much their partner could be willing to invest later on during the breeding attempt. This may be critical for individuals to estimate and to adjust their own effort. In the future, finer-scale analyses may also allow us to understand whether and how individuals respond strategically to each other’s behaviour, for example by taking turns (cf. Johnstone & Savage, 2019; Savage et al., 2017). Given growing evidence that nest building improves with experience (Muth & Healy, 2011), it is also important to establish whether pairs learn and refine their cooperative nest building strategies over time. Although there was little evidence that age was an important factor in our analyses, future work will be vital to determine whether and how the prior history of specific partners shapes their behaviour and reproductive success. Finally, investigations of nest building may also contribute to our understanding of animal architecture. To date, the majority of research on cooperative architecture has focused on the nests and mounds of eusocial insects, where the colony is the unit of reproduction. Cooperative nest-building in birds may provide useful opportunities to understand how variation in conflicts of interest influences the adaptive value of cooperating to build structures for mutual benefit, and the proximate mechanisms through which this is achieved.

## Acknowledgements

We are very grateful to the Gluyas family and staff at Pencoose Farm as well as to the people of Stithians who allowed us to carry out fieldwork. We thank Emily Cuff for collecting some of the video data in the field. This research was supported by a BBSRC David Phillips Fellowship (BB/H021817/2) and a Human Frontier Science Program grant (RG0049/2017) to A.T. R.H. was supported by a Natural Environment Research Council GW4 studentship (NERC 107672G).

## Appendix

## i. Behavioural variables

**Table A 1.** Behaviours (state events) jackdaws showed in their nest box (N = 62 observations; N = 60 for chatter). The second and third column summarise the percentage of time pairs showed each behaviour. The subsequent two columns indicate the amount of time (percentage of observation length) both sexes exhibited a particular behaviour. The behaviours of the sexes do not always add up to the cumulative amount because in some instances a bird was not identifiable.

| Behaviour | Percent of time<br>(out of obs.<br>length)<br>Mean ± SD | Percent of time<br>(out of time<br>in box)<br>Mean ± SD | Females ♀:<br>percent of time<br>(obs. length)<br>Mean ± SD | Males ♂:<br>percent of time<br>(obs. length)<br>Mean ± SD |
| --- | --- | --- | --- | --- |
| In box (cum.) | 37.96 ± 30.50 | - | 16.06 ± 15.99 | 15.50 ± 15.04 |
| Box occupied | 29.09 ± 19.64 | - | - | - |
| In box (both) | 8.99 ± 13.56 | 23.17 ± 24.58 | - | - |
| Vigilance | 15.73 ± 19.24 | 37.83 ± 19.32 | 5.58 ± 8.69 | 7.57 ± 10.47 |
| Build nest | 9.59 ± 8.39 | 27.34 ± 16.92 | 4.66 ± 4.95 | 3.05 ± 3.48 |
| Form cup | 1.67 ± 1.70 | 4.71 ± 4.01 | 0.74 ± 0.90 | 0.60 ± 0.68 |
| Modify | 0.05 ± 0.13 | 0.15 ± 0.52 | 0.024 ± 0.075 | 0.003 ± 0.014 |
| Chatter | 0.92 ± 1.70 | 1.93 ± 2.50 | 0.33 ± 0.80 | 0.33 ± 0.77 |

**Table A 2.**
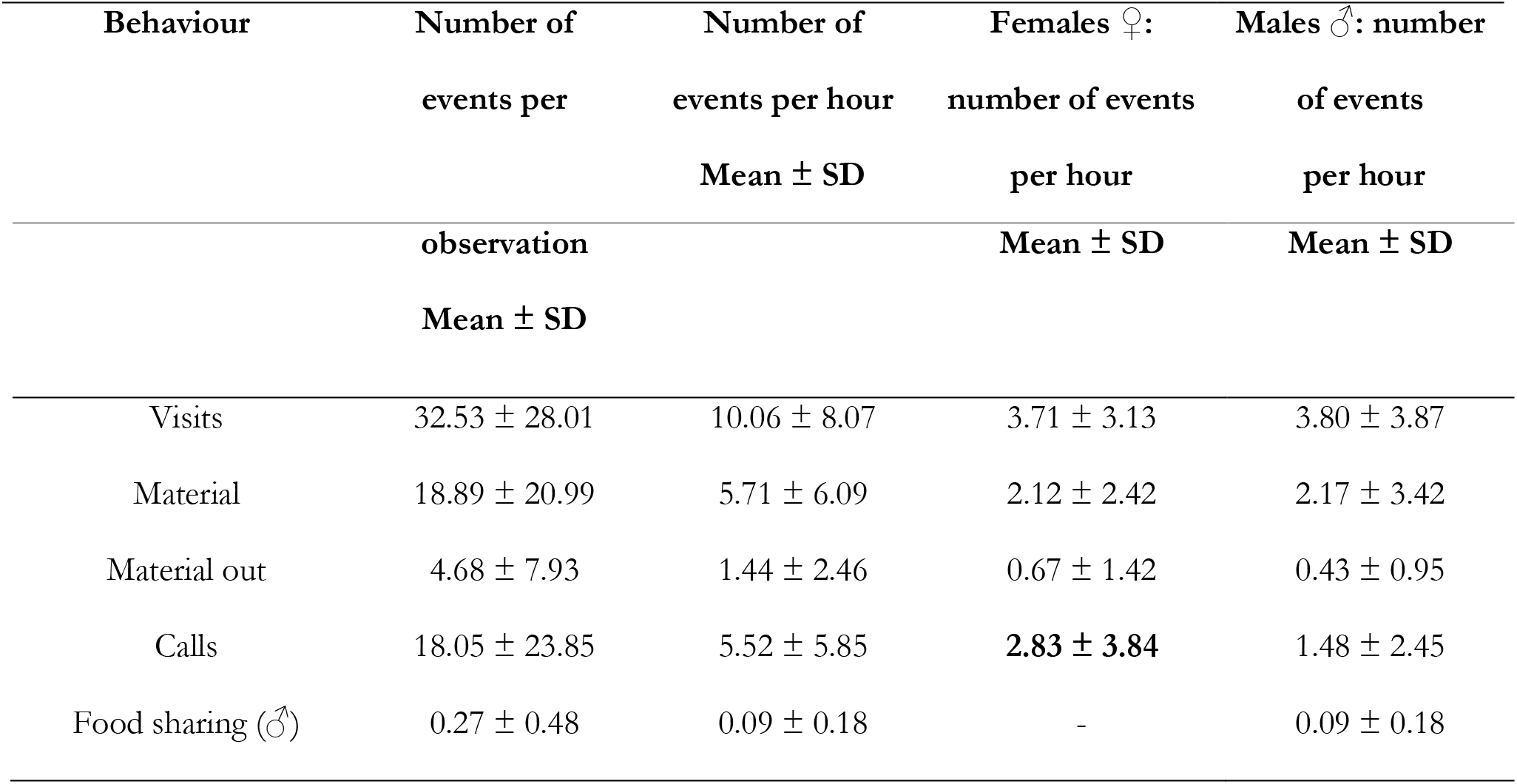
Behaviours (point events) shown by jackdaw pairs (N = 62 observations). The second and third column describe the total number of events per observation and per hour, respectively. The last two columns summarise the number of events per hour for both sexes separately. Please note the behaviours of the sexes do not add up to the cumulative amount, as individuals were sometimes unidentifiable.

**Table A 3.** Relationship between behaviours shown by jackdaws and age.

| Response | Fixed effects | $\beta$ | SE ($\beta$) | $X^2$ | df | 95 % CI | 95 % CI | *P* |
| --- | --- | --- | --- | --- | --- | --- | --- | --- |
| variable |  |  |  |  |  | (lower) | (upper) |  |
| (model) |  |  |  |  |  |  |  |  |
| In box | Age | 0.091 | 0.112 | 0.398 | 1 | - 0.127 | 0.310 | 0.528 |
| (LMM) | Sex * Age | - 0.059 | 0.116 | 0.264 | 1 | - 0.285 | 0.165 | 0.607 |
| Build nest | Age | 0.054 | 0.079 | 0.573 | 1 | - 0.100 | 0.208 | 0.449 |
| (LMM) | Sex * Age | - 0.011 | 0.080 | 0.017 | 1 | - 0.166 | 0.145 | 0.895 |
| Vigilance | Age | 0.050 | 0.099 | 0.036 | 1 | - 0.140 | 0.241 | 0.849 |
| (LMM) | Sex * Age | - 0.060 | 0.099 | 0.367 | 1 | - 0.251 | 0.132 | 0.545 |
| Build vs. | Age | 0.027 | 0.059 | 0.205 | 1 | - 0.088 | 0.141 | 0.651 |
| vigilance |  |  |  |  |  |  |  |  |
| (LMM) |  |  |  |  |  |  |  |  |
| Visits | Age | 0.165 | 0.081 | 2.380 | 1 | 0.005 | 0.324 | 0.123 |
| (GLMM) | Sex * Age | - 0.111 | 0.085 | 1.712 | 1 | - 0.278 | 0.055 | 0.191 |
| Material out | Age | 0.222 | 0.243 | 0.275 | 1 | - 0.255 | 0.698 | 0.600 |
| (GLMM) | Sex * Age | - 0.247 | 0.322 | 0.590 | 1 | - 0.878 | 0.383 | 0.442 |
| Form cup | Age | 0.048 | 0.037 | 1.380 | 1 | - 0.025 | 0.120 | 0.240 |
| (LMM) | Sex * Age | - 0.022 | 0.038 | 0.347 | 1 | - 0.096 | 0.051 | 0.556 |
| Chatter | Age | 0.059 | 0.032 | 2.112 | 1 | - 0.002 | 0.121 | 0.146 |
| (LMM) | Sex * Age | - 0.043 | 0.036 | 1.389 | 1 | - 0.113 | 0.027 | 0.239 |
| Calls | Age | - 0.044 | 0.148 | 1.046 | 1 | - 0.334 | 0.247 | 0.306 |
| (GLMM) | Sex * Age | 0.280 | 0.169 | 2.378 | 1 | - 0.051 | 0.611 | 0.098 |

**Figure A 1.**
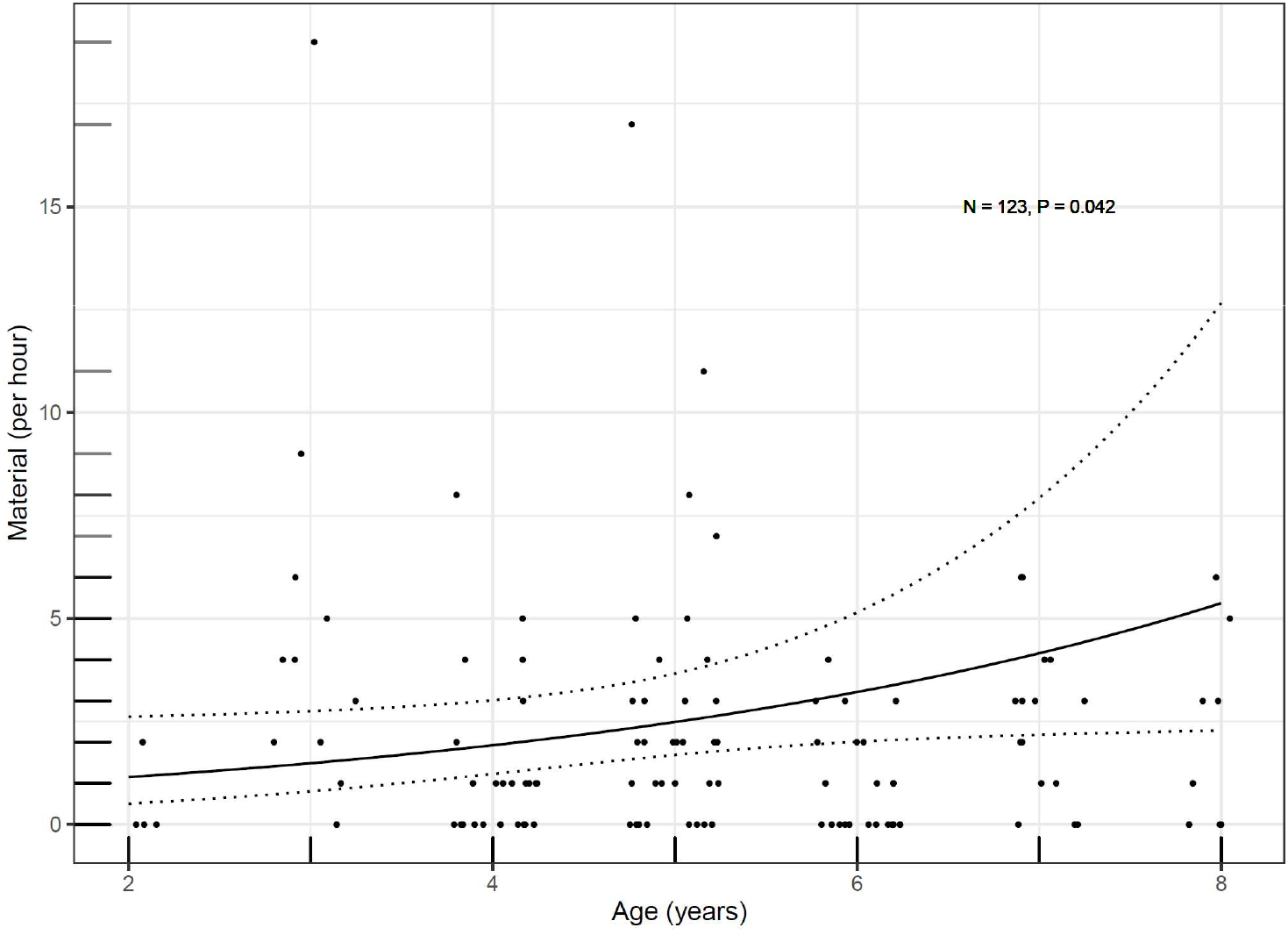
The number of material brought by individual birds (rounded, per hour) plotted against their age (horizontal jitter used to make datapoints more distinguishable). The relationship was non-significant when removing four individuals that were two years old (*P* = 0.200). The continuous fitted line corresponds to the model output; dotted lines show the 95 % confidence intervals.

## ii. Fitness correlates: Principal Component Analyses (PCA)

**Table A 4.** Loadings of behavioural variables onto the first two principal components of the PCA_All_ including nine different behaviours (N = 59).

| Behaviour | PC1 | PC2 |
| --- | --- | --- |
| In box (cumulative) | - 0.456 | 0.175 |
| In box (both birds) | - 0.354 | 0.391 |
| Build nest | - 0.344 | - 0.385 |
| Material | - 0.309 | - 0.449 |
| Material out | - 0.224 | - 0.478 |
| Form cup | - 0.266 | - 0.226 |
| Vigilance | - 0.380 | 0.390 |
| Chatter | - 0.297 | 0.048 |
| Calls | - 0.317 | 0.180 |
| Variation explained | 46.54 | 25.22 |

**Table A 5.** Loadings of behavioural variables related to nest building onto PC1Efffort and PC2Efffort of the PCAEfffort (N = 61).

| Behaviour | PC1 | PC2 |
| --- | --- | --- |
| Build nest | - 0.541 | 0.017 |
| Material | - 0.543 | - 0.123 |
| Material out | - 0.496 | - 0.558 |
| Form cup | - 0.408 | 0.821 |
| Variation explained | 69.18 | 18.86 |

**Table A 6.** Loadings of behavioural variables (relative female contribution) onto PC1 and PC2 of the PCA_DoL_ (N = 61).

| Behaviour | PC1 | PC2 |
| --- | --- | --- |
| Build nest ♀ | 0.700 | - 0.118 |
| Material ♀ | 0.116 | 0.992 |
| Vigilance ♀ | - 0.704 | 0.046 |
| Variation explained | 58.16 | 33.06 |

## iii. Fitness correlates: mixed models

**Table A 7.** Different models to examine the effect of behaviours on proxies for reproductive success. In all models we accounted for the ‘year’, ‘day’, ‘food sharing’ and ‘female age’ as fixed effects. ‘Pair ID’ and ‘site’ were included as random effects in all models. The column ‘best model’ shows which model had the lowest AICc in cases where we performed model selection. The last two columns show the estimate, standard error, and P-value for the instances where we did not use AICc but constructed single models.

| Response variable | Concept (models) | Fixed effects | Error structure | Best model ($\Delta$ AICc) | $\beta \pm SE (\beta)$ | *P* |
| --- | --- | --- | --- | --- | --- | --- |
| Relative lay date | PCA<sub>All</sub> (5) | PC1<sub>All</sub>, PC2<sub>All</sub> | GLMM: | Year, Day (3.3) | - | - |
| | PCA<sub>Effort</sub> (1) | PC1<sub>Effort</sub> | COM- | - | - 0.012 ± 0.012 | 0.286 |
|  | **PCA<sub>DoL</sub> (6)** | PC1<sub>DoL</sub>, Material by ♀ | Poisson | Material ♀ (4.7) | - | - |
| | Synchrony (1) | Together in box | | - | 0.002 ± 0.003 | 0.546 |
| Clutch size | PCA<sub>All</sub> (5) | PC1<sub>All</sub>, PC2<sub>All</sub> | GLMM: | Null model (0.0) | - | - |
| | PCA<sub>Effort</sub> (1) | PC1<sub>Effort</sub> | COM- | - | 0.005 ± 0.017 | 0.753 |
|  | PCA<sub>DoL</sub> (6) | PC1<sub>DoL</sub>, Material by ♀ | Poisson | Null model (3.6) | - | - |
| | Synchrony (1) | Together in box | | - | - 0.002 ± 0.002 | 0.424 |
| Volume of 1<sup>st</sup> egg | PCA<sub>All</sub> (5) | PC1<sub>All</sub>, PC2<sub>All</sub> | LMM: | Null model (3.2) | - | - |
| | PCA<sub>Effort</sub> (1) | PC1<sub>Effort</sub> | Gaussian | - | - 0.032 ± 0.071 | 0.656 |
|  | PCA<sub>DoL</sub> (6) | PC1<sub>DoL</sub>, Material by ♀ |  | Null model (3.4) | - | - |
| | **Synchrony (1)** | Together in box | | - | - 0.018 ± 0.008 | **0.019** |
| Volume of 3<sup>rd</sup> egg | PCA<sub>All</sub> (5) | PC1<sub>All</sub>, PC2<sub>All</sub> | LMM: | Null model (9.8) | - | - |
| | PCA<sub>Effort</sub> (1) | PC1<sub>Effort</sub> | Gaussian | - | - 0.063 ± 0.087 | 0.469 |
|  | PCA<sub>DoL</sub> (6) | PC1<sub>DoL</sub>, Material by ♀ |  | Null model (8.5) | - | - |
| | **Synchrony (1)** | Together in box | | - | - 0.026 ± 0.010 | **0.007** |

## iv. Behaviour and fitness proxies: Repeatability

**Table A 8.** Repeatability estimates for different behaviours of 16 pairs that were measured repeatedly. State events (SE) were Box-Cox transformed to approximate assumptions for Gaussian data.

| Behaviour | Repeatability | SE | 2.5 % CI | 97.5 % CI | *P* |
| --- | --- | --- | --- | --- | --- |
| In box (cumulative) (SE) | 0 | 0.152 | 0 | 0.490 | 1 |
| In box (both birds) (SE) | 0 | 0.155 | 0 | 0.518 | 1 |
| Vigilance (SE) | 0 | 0.154 | 0 | 0.509 | 1 |
| Nest build (SE) | 0.094 | 0.176 | 0 | 0.581 | 0.378 |
| Material | 0 | 0.131 | 0 | 0.437 | 1 |
| Material out | 0.031 | 0.181 | 0 | 0.639 | 0.389 |
| Form cup (SE) | 0 | 0.146 | 0 | 0.475 | 1 |
| Chatter (SE) | 0 | 0.152 | 0 | 0.494 | 0.5 |
| Calls | 0 | 0.103 | 0 | 0.357 | 1 |
| Food sharing | Not converged | - | - | - | - |

**Table A 9.**
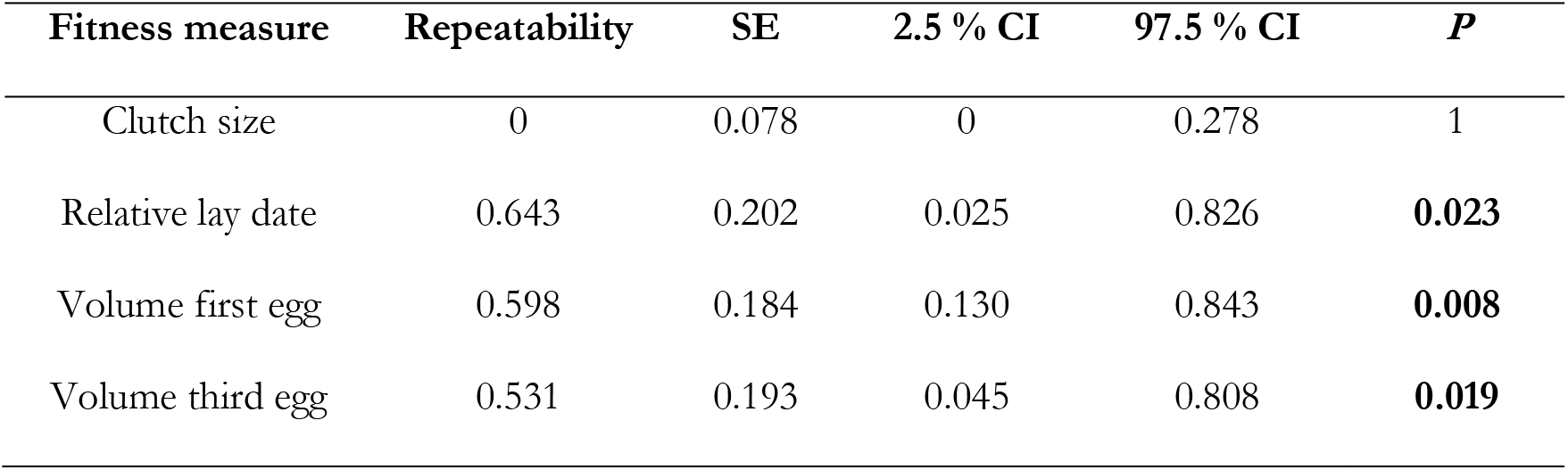
Repeatability estimates for different correlates of reproductive success.

